# Tip60 protects against amyloid-β peptide-induced transcriptomic alterations via different modes of action in early versus late stages of neurodegenerative progression

**DOI:** 10.1101/2020.06.09.142885

**Authors:** Haolin Zhang, Bhanu Chandra Karisetty, Akanksha Bhatnagar, Ellen M. Armour, Mariah Beaver, Tiffany V. Roach, Sina Mortazavi, Shreya Mandloi, Felice Elefant

## Abstract

Alzheimer’s disease (AD) is an age-related neurodegenerative disorder hallmarked by amyloid-β (Aβ) plaque accumulation, neuronal cell death, and cognitive deficits that worsen during disease progression. Histone acetylation dysregulation, caused by an imbalance between reduced histone acetyltransferases (HAT) Tip60 and increased histone deacetylase 2 (HDAC2) levels, can directly contribute to AD pathology. However, whether such AD-associated neuroepigenetic alterations occur in response to Aβ peptide production and can be protected against by increasing Tip60 levels over the course of neurodegenerative progression remains unknown. Here we profile Tip60 HAT/HDAC2 dynamics and transcriptome-wide changes across early and late stage AD pathology in the *Drosophila* brain produced solely by human amyloid-β_42_. We show that early Aβ_42_ induction leads to disruption of Tip60 HAT/HDAC2 balance during early neurodegenerative stages preceding Aβ plaque accumulation that persists into late AD stages. Correlative transcriptome-wide studies reveal alterations in biological processes we classified as transient (early-stage only), late-onset (late-stage only), and constant (both). Increasing Tip60 HAT levels in the Aβ_42_ fly brain protects against AD functional pathologies that include Aβ plaque accumulation, neural cell death, cognitive deficits, and shorter life-span. Strikingly, Tip60 protects against Aβ_42_-induced transcriptomic alterations *via* distinct mechanisms during early and late stages of neurodegeneration. Our findings reveal distinct modes of neuroepigenetic gene changes and Tip60 neuroprotection in early versus late stages in AD that can serve as early biomarkers for AD, and support the therapeutic potential of Tip60 over the course of AD progression.

## INTRODUCTION

Alzheimer’s Disease (AD) is a debilitating age-associated neurodegenerative disorder (ND) characterized by amyloid-β (Aβ) plaque accumulation, neural cell death, and cognitive decline that worsens as the disease advances. The progression of AD is a complex interaction between age, genetic and environmental factors (Karch et al. 2014; Masters et al. 2015; Sanchez-Mut and Graff 2015) in which epigenetic gene regulatory mechanism is believed to play a critical role. Compelling evidence by our group and others demonstrates that neural histone acetylation dysregulation, caused by an imbalance between histone acetyltransferases (HATs) and histone deacetylases (HDACs), can directly contribute to AD pathology (Saha and Pahan 2006; Sanchez-Mut and Graff 2015). Downregulation of the HAT Tip60 (KAT5) (Panikker et al. 2018) and upregulation of HDAC2 (Graff et al. 2012) has been associated with epigenetic repression of critical neuroplasticity genes in AD animal models and patients. Restoring such alterations in Tip60/HDAC2 balance has been shown to protect against AD-associated pathologies in an AD *Drosophila* model expressing amyloid precursor protein (APP) (Panikker et al. 2018).

Questions with significant clinical relevance that are critical to address are whether such AD-associated neuroepigenetic alterations occur in response to Aβ production, are constant or dynamic during neurodegenerative progression, and can be protected against by increasing Tip60 over the course of the disease. If the neuroepigenome and corresponding transcriptome are highly dynamic during AD progression, elucidating these changes would aid in the identification of early AD biomarkers and specific therapeutic targets for a particular disease stage. Recent studies carrying out genome-wide profiling of epigenetic modification changes in the CK-p25 AD-associated mouse model revealed transient, consistent and late alterations in histone acetylation domains and concomitant gene expression profiles (Gjoneska et al. 2015). These studies support a model in which an AD early-response epigenome and AD late-response epigenome represent different stages of disease progression. Nevertheless, the impact of restoring Tip60 HAT levels on epigenetic gene dysregulation during early and late stages of neurodegeneration progression remain unknown, given the inaccessibility of human brain samples from young patients exhibiting mild AD pathologies.

To address this need, here we profile Tip60 HAT/HDAC2 dynamics and transcriptional changes across early and late stage AD pathology in the *Drosophila* brain produced solely by induction of neurotoxic human Aβ_42_. We show that early Aβ_42_ induction leads to disruption of Tip60 HAT/HDAC2 balance that occurs during early neurodegeneration preceding Aβ plaque accumulation that persists into later AD stages. Transcriptome-wide studies reveal alterations in biological processes we classified as transient (early-stage only), late-onset (late-stage only), and constant (both). Strikingly, increased Tip60 in the brain protects against Aβ_42_-induced transcriptome-wide alterations *via* distinct mechanisms during early and state stages of neurodegeneration, suggesting greater specificity for Tip60 protection against Aβ_42_-induced alterations during earlier stages of the disease. Our findings provide new insights into distinct modes of epigenetic gene changes and Tip60 neuroprotection in early versus late stages of AD that can serve as early biomarkers for AD, and support the therapeutic potential of Tip60 over the course of AD progression.

## RESULTS

### Tip60 protects against Aβ_42_-induced amyloid-β plaque accumulation and apoptotic driven neurodegeneration in the *Drosophila* brain

We first asked whether Tip60 HAT action plays a neuroprotective role in neuropathology produced solely by induction of neurotoxic amyloid-β in the nervous system. To achieve this, we used the well-characterized AD fly model that expresses a secreted Aβ_42_ fusion protein under control of the UAS promoter. Pan-neuronal driven expression of the Aβ_42_ construct by the elav-GAL4 driver has been shown to cause AD-associated neuropathology that includes the formation of age-dependent diffused amyloid deposits in the brain, age-dependent learning defects, and extensive neurodegeneration (Iijima et al. 2004). We generated a double transgenic Aβ_42_;Tip60 fly line that enables induction of increased Tip60 HAT levels specifically in the nervous system in this Aβ_42_-induced neurotoxicity background. Validation of transgene expression in the brain confirmed similar levels of Aβ_42_ between Aβ_42_ and Aβ_42_;Tip60 fly lines and that Tip60 levels are effectively increased in the brain when induced by the pan-neuronal elav-GAL driver in our Aβ_42_;Tip60 fly model (Supplemental Fig. S1A and S1B).

We first assessed whether increasing Tip60 HAT levels in the brain could protect against the amyloid plaque deposits that accumulate in the aged 28-day old fly brain as a result of Aβ_42_ induction. We focused our studies on the mushroom body (MB) Kenyon cell region of the brain as we have shown that Tip60 is robustly expressed in the MB and is critical for MB function in learning and memory (Xu et al. 2014). As previously documented, anti-Aβ immunofluorescence studies (Iijima et al. 2004) revealed that Aβ_42_ expression in the *Drosophila* brain results in diffuse amyloid deposits that do not appear in abundance in the MB until flies are aged to 28 days (Fig. 1Aii and 1Bii). These Aβ plaque deposits are unobservable in an earlier AD stage modeled in third-instar larvae (Fig. 1Ai and 1Bi). Strikingly, increasing Tip60 levels in the Aβ_42_ background shows a reduction in both the number and size of Aβ plaques (Fig. 1Aii and 1Bii).

**Figure 1.**
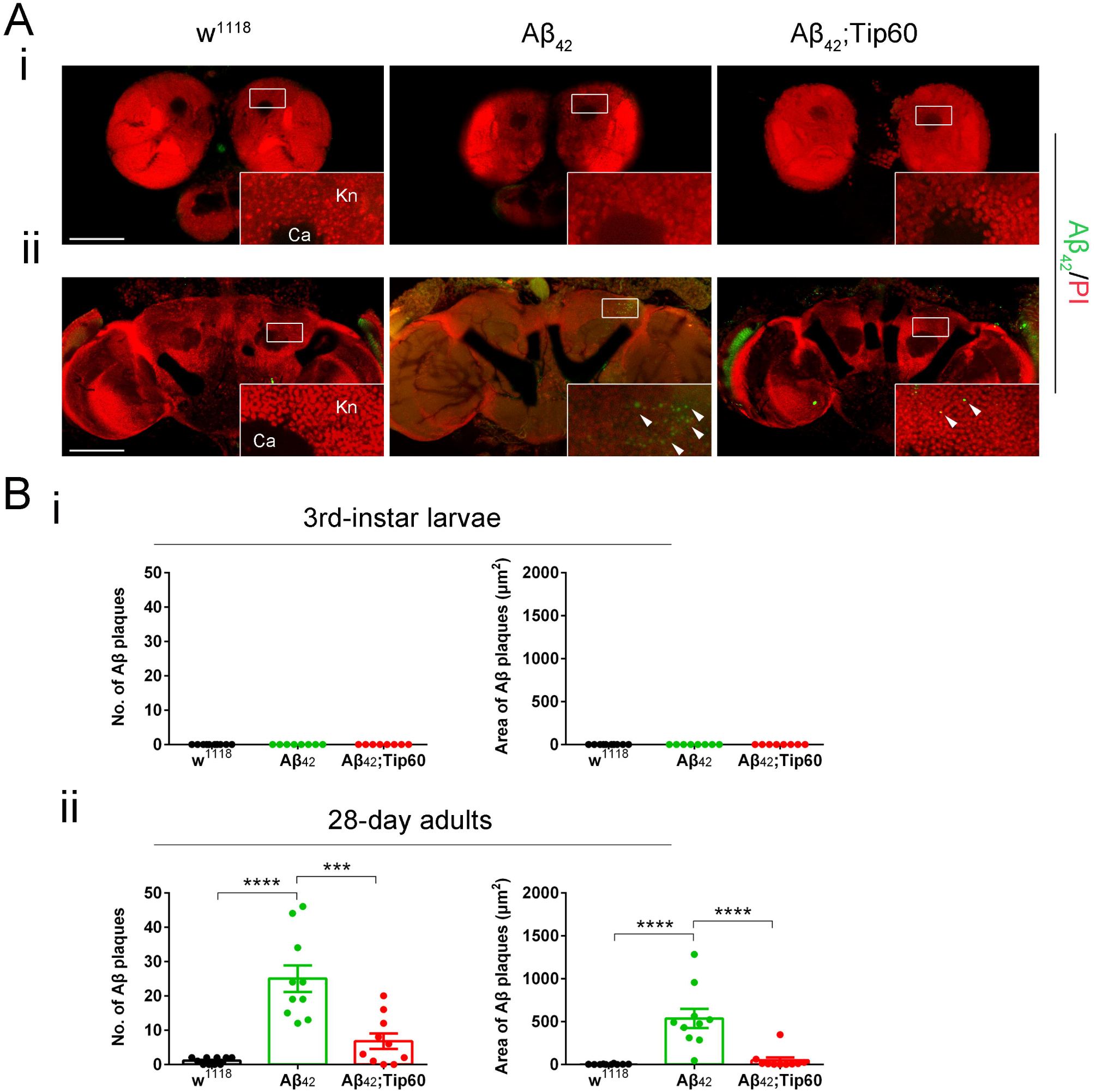
Increased Tip60 levels in the Aβ_42_ brain protects against Aβ plaque accumulation in the fly brain mushroom body Kenyon cell (Kn) region. (A) Representative images in each genotype. Aβ plaques were stained with anti-Aβ_42_ antibody (green). Nuclei were stained with PI (red). The Kn cell region (boxed) was zoomed in to display Kn cells and Aβ plaques. (i) Immunostaining of brains of 3rd-instar larvae shows no Aβ_42_ signal in all three genotypes. (ii) Immunostaining of brains of 28-day adults shows evident Aβ plaques in Aβ_42_ flies compared to w^1118^ flies. Overexpression of Tip60 in the Aβ_42_ background reduces Aβ plaques. Arrowheads indicate Aβ plaques. No Aβ_42_ signal was detected in the Calyx (Ca) region. Scale bar represents 100 μm. (B) Aβ plaque was quantified by both number and size. (i) Quantification of Aβ plaque numbers and areas in the 3rd-instar larval brain Kn region. (ii) Quantification of Aβ plaque numbers and area in the 28-day adult brain Kn region. n = 8 ∼ 10. \*\*\**p* < 0.001, \*\*\*\**p* < 0.0001; one-way ANOVA with Tukey’s multiple comparisons test. All data are shown as mean ± s.e.m.

We next asked whether increased Tip60 also protects against Aβ_42_-induced neuronal apoptosis by examining the MB Kenyon cell region in the brain using terminal deoxynucleotidyl transferase dUTP nick end labeling (TUNEL). Control experiments using DNase LJ treated brains confirm apoptotic TUNEL specificity (Supplemental Fig. S2A and S2B). TUNEL staining in the third-instar larval brain revealed a minimal of apoptotic cells in this early Aβ_42_-induced neurodegeneration model that was comparable to wild type control brains (Fig. 2Ai and 2Bi). In contrast, aged 28-day flies display a significantly higher level of apoptotic neuronal cell death when compared to control wild-type brains (Fig. 2Aii and 2Bii). Increasing Tip60 levels shows a drastic reduction in the number of apoptotic cells (Fig. 2Aii and 2Bii). Taken together, our results demonstrate that the accumulation of both Aβ_42_-induced amyloid plaques and neuronal apoptotic cell death in the Kenyon cell region of the MB does not occur until later stages of AD-associated neuropathology and can be effectively protected against by increasing Tip60 levels in the brain.

**Figure 2.**
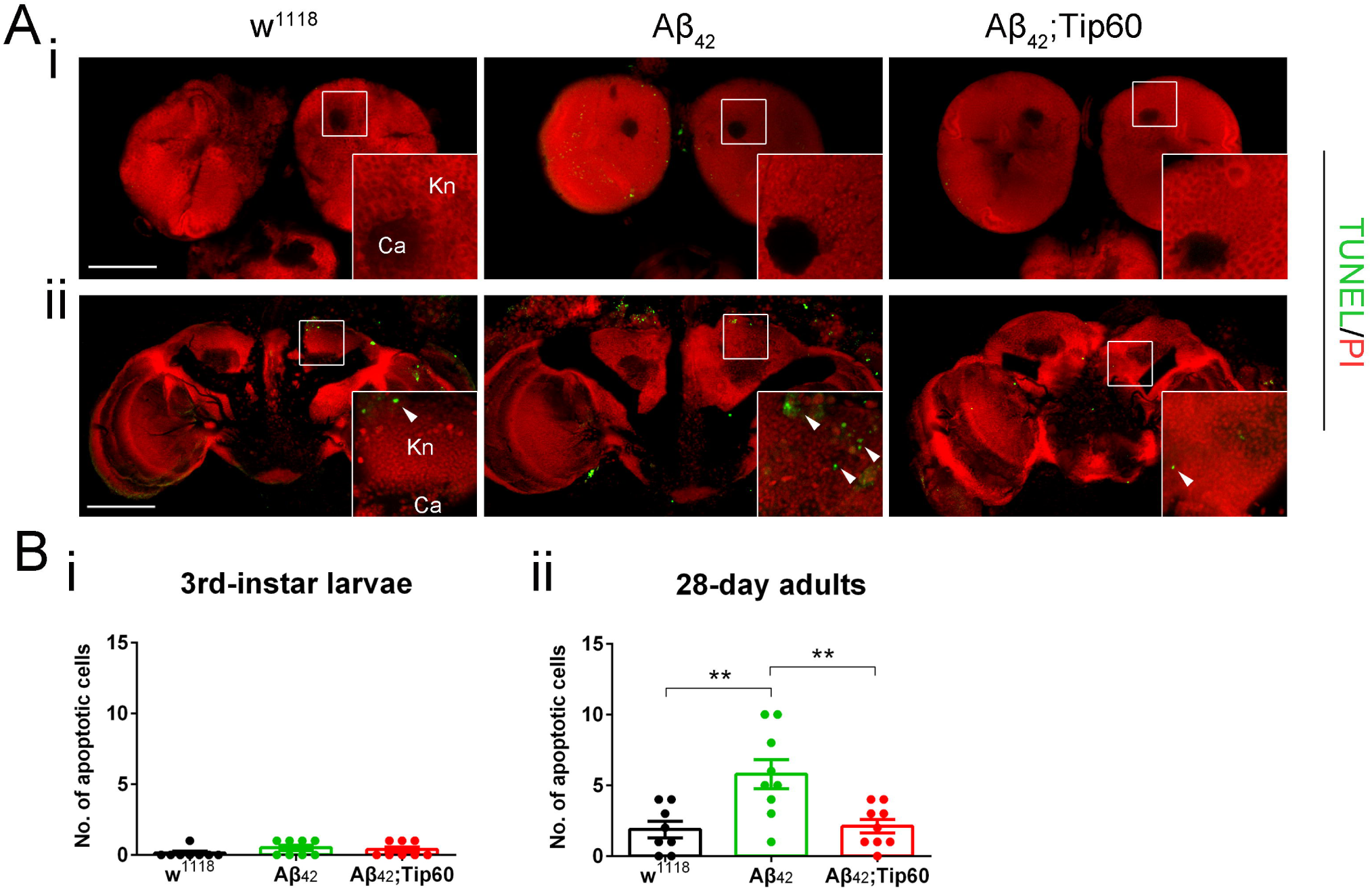
Increased Tip60 levels in the Aβ_42_ brain protects against apoptosis in the fly brain mushroom body Kenyon cell (Kn) region. (A) Representative confocal images of neuronal apoptosis visualized by TUNEL staining (green) of brains expressing indicated transgenes driven by pan-neuronal driver elav-Gal4. Nuclei were stained with PI (red). The Kn cell region (boxed) was zoomed in to display Kn cells and apoptotic cells. (i) Immunostaining of brains of 3rd-instar larvae shows minimal apoptotic signals in all three genotypes. (ii) Immunostaining of brains of 28-day adults shows evident apoptosis in Aβ_42_ flies compared to w^1118^ flies. Overexpression of Tip60 in the Aβ_42_ background rescues the apoptosis phenotype. Arrowheads indicate TUNEL positive apoptotic cells. No apoptotic signal was detected in the Calyx (Ca) region. Scale bar represents 100 μm. (B) Apoptosis was quantified by the number of apoptotic cells. (i) Apoptosis quantification of 3rd-instar larval brain Kn region. (ii) Apoptosis quantification of 28-day adult brain Kn region. n = 8 ∼ 9. \*\**p* < 0.01; one-way ANOVA with Tukey’s multiple comparisons test. All data are shown as mean ± s.e.m.

### Tip60 HAT action protects against Aβ_42_-induced deficits in learning and memory, locomotion activity, and longevity

Thus far, we demonstrated that Tip60 protects against Aβ_42_-induced human conserved AD neuropathology in the fly brain at a cellular level. Additional human AD pathologies feature learning and short-term memory (STM) loss, movement deficits, and early lethality. Each of these functional deficits is conserved in the AD-associated aged Aβ_42_ model and has been previously documented and quantified (Iijima et al. 2004; Iijima-Ando et al. 2008; Wu et al. 2019). Thus, we asked whether these deficits occur during the early stages of Aβ_42_-induced neuropathology and whether increasing Tip60 levels can protect against these deficits throughout neurodegenerative progression.

To assess learning and memory deficits during early AD neurodegenerative progression modeled in Aβ_42_ larvae, we carried out a single odor paradigm for olfactory associative learning and STM (Honjo and Furukubo-Tokunaga 2005). Larvae expressing either Aβ_42_, Aβ_42_;Tip60 or w^1118^ wild-type control under the control of the pan-neuronal elav-GAL4 driver were first conditioned to associate the given odor, linalool (LIN), to an appetitive gustatory reinforcer, sucrose (SUC). These larvae were exposed to LIN for 30 minutes on an agar plate containing SUC. After conditioning, the larvae were tested for an olfactory response on the test plate (Supplemental Fig. S3Ai). Control experiments show equivalent sensory acuities to odor and natural reinforcer, confirming the validity of the assay (Supplemental Fig. S3Aiii and S3Aiv). Response index (RI) of each genotype were normalized using their respective locomotion speed for equivalent comparison (Supplemental Fig. S3Av). Larval response after 0 minutes and 30 minutes of LIN/SUC conditioning were compared to test learning and STM, respectively. In comparison to the control larvae that had been exposed to LIN/distilled water (DW), LIN/SUC conditioned w^1118^ showed enhanced migration, and high ΔRI at 0 minutes (Supplemental Fig. S3Aii) and 30 minutes (Fig. 3Ai), indicating normal learning and STM. Aβ_42_ larvae showed significantly lower ΔRI at both 0 minutes (Supplemental Fig. S3Aii) and 30 minutes (Fig. 3Ai), indicating defects in both learning and STM. Increasing Tip60 HAT levels in the nervous system in Aβ_42_;Tip60 larvae protected against these Aβ_42_-induced learning deficits (Supplemental Fig. S3Aii) as well as elicited STM (Fig. 3Ai) under Aβ_42_-induced early neurodegenerative conditions.

**Figure 3.**
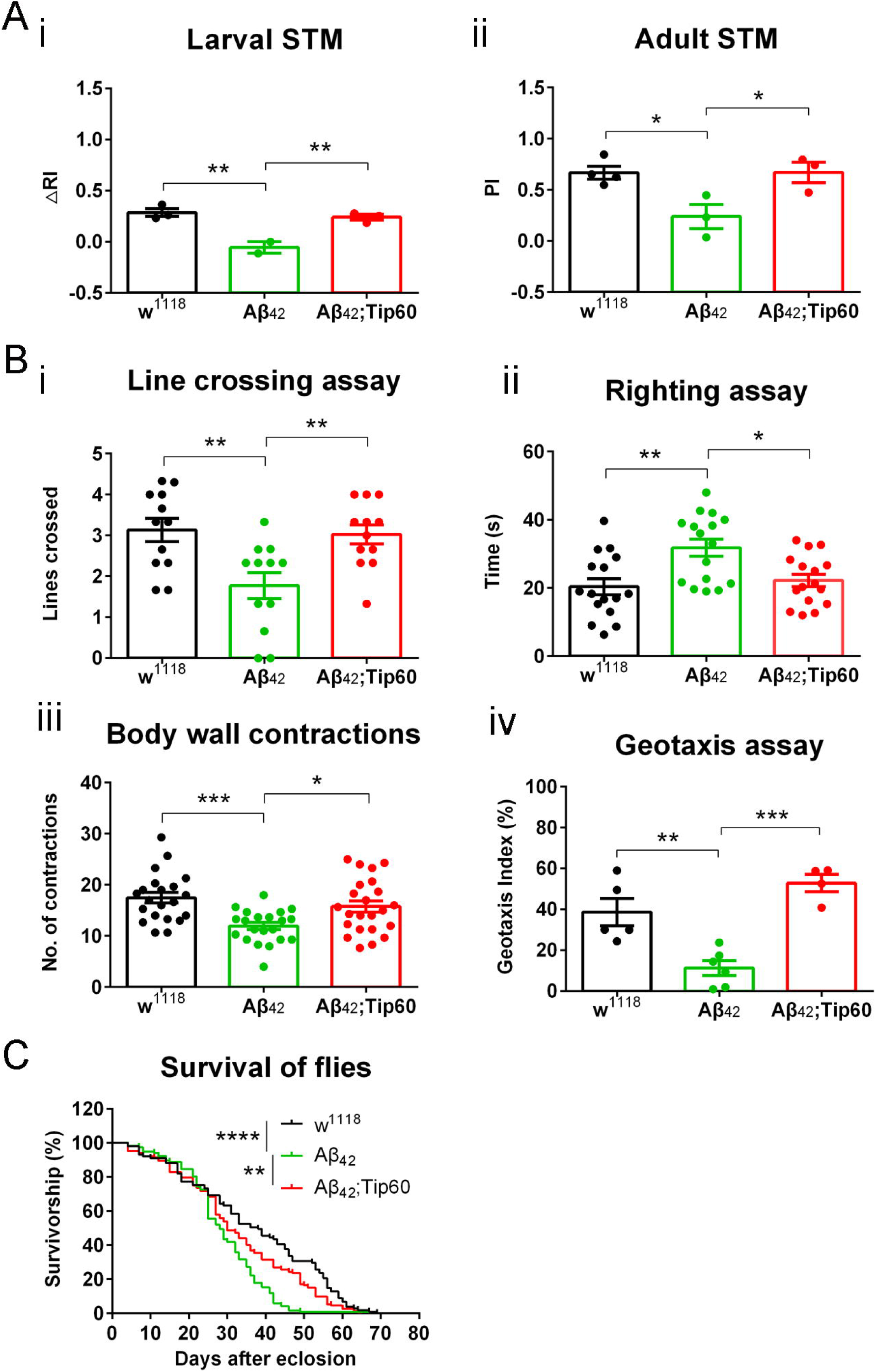
Increasing Tip60 levels in the Aβ_42_ fly brain protects against AD associated pathologies. (A) Tip60 restoration rescues the STM defects induced by Aβ pathology. (i) Tip60 restoration improves STM performance in Aβ_42_ neurodegenerative larvae. n = 2 ∼ 3. Each biological repeat has 60 ∼ 100 larvae. (ii) Tip60 restoration improves STM performance in Aβ_42_ neurodegenerative adults. n = 3 ∼ 4. Each biological repeat has 30 ∼ 60 adult flies. (B) Aβ_42_ fly locomotion deficits are effectively rescued by Tip60 overexpression. (i) Larval line crossing assay. n = 12. (ii) Larval righting assay. n = 16. (iii) Larval body wall contraction assay. n = 21 ∼ 23. (iv) Adult negative geotaxis assay. n = 4 ∼ 6. Each biological repeat has at least 30 flies. (C) The expression of Aβ_42_ causes a shorter life-span that is partially rescued by Tip60. n = 100 ∼ 160. \**p* < 0.05, \*\**p* < 0.01, \*\*\**p* < 0.001, \*\*\*\**p* < 0.0001; one-way ANOVA with Tukey’s multiple comparisons test (STM and locomotion assays); Log-rank test with multiple adjustments (survival assay). All data except survival curve are shown as mean ± s.e.m.

To assess learning and memory in aged AD flies, we used the well-established *Drosophila* adult olfactory shock learning and memory assay (Malik and Hodge 2014). Flies aged to 28 days and expressing either Aβ_42_, Aβ_42_;Tip60, or w1118 wild-type control under the control of the pan-neuronal elav-GAL4 driver were used for these studies. The adult flies were presented the first odor (MCH or OCT) that was paired with a 65 V shock for 60 seconds, followed by a 30 seconds rest period without odor or shock. The flies were then presented the second odor (OCT or MCH) without shock, followed by a rest period without odor or shock. The two odors were used in a counterbalanced design, with half of the flies used in the calculation of the performance index (PI) being trained to OCT-shock and the other half to MCH-shock. Flies were removed from the T-maze, allowed to rest for 30 minutes, and moved back to the T-maze for the STM test by choosing between the two scents. A higher avoidance of the shock-paired odor indicates a better memory performance, which is presented by a higher PI. As expected, after 30 minutes, w^1118^ control flies showed a PI of 0.67 (Fig. 3Aii), which is indicative of significant shock-paired odor avoidance and effective STM. In contrast, after 30 minutes, Aβ_42_ flies showed a marked reduced PI of 0.24 (Fig. 3Aii), which is indicative of substantial deficits in STM reflected. Strikingly, Aβ_42_;Tip60 flies showed a PI of 0.67 (Fig. 3Aii), which is comparable to that of control w^1118^ flies, indicating that increased levels of Tip60 protect against STM deficits that persist into late stages of Aβ_42_-induced neurodegeneration. Control experiments revealed similar olfactory acuities and electric shock avoidance between all genotypes (Aβ_42_ and Aβ_42_;Tip60 flies are similar to control w^1118^) (Supplemental Fig. S3Bi and S3Bii), indicating that these functions themselves are not compromised by Aβ_42_ induction.

To assess whether motor neuron function is compromised during early stages of Aβ_42_-induced neurodegeneration, a series of functional locomotion assays that include locomotion, contraction, and righting assays were performed as previously described (Mudher et al. 2004). Larvae expressing either Aβ_42_, Aβ_42_;Tip60 or w^1118^ wild-type control under the control of the pan-neuronal elav-GAL4 driver were used for each of these assays. Contraction capability, locomotion function, and the ability to perform a complex motor task such as righting provide direct quantifiable evidence of motor neuron function. Our studies revealed that in each of these larval motor neuron function assays, the Aβ_42_ larval function was significantly impaired relative to the control w^1118^ larvae (Fig. 3Bi-iii). In contrast, Aβ_42_;Tip60 larvae displayed motor neuron function similar to control w^1118^ larvae (Fig. 3Bi-iii), indicating that increasing Tip60 levels protects against early Aβ_42_ functional motor deficits. To test whether Tip60 neuroprotection persists into later stages of neurodegeneration, we assessed motor neuron function in aged 28-day adult flies using the well-established geotaxis assay (Krashes and Waddell 2008). Negative geotaxis assay vials were positioned vertically, and the geotaxis index was scored as the percentage of flies that could crawl to the top tube after 10 seconds (Krashes and Waddell 2008). The results revealed that Aβ_42_ flies showed reduced climbing performance, whereas the performance of Aβ;Tip60 flies was efficiently improved (Figure 3Biv). These results suggest Tip60-mediated protection against early motor deficits persists into later stages of dysfunction.

We next assessed the effects of increasing Tip60 levels on Aβ_42_ fly life-span. Results from this longevity assay revealed that 50% of Aβ_42_ flies were alive at 28 days as compared to approximately 70% for w^1118^ and Aβ_42_;Tip60 flies (Fig. 3C). Further, after 50 days, there were no Aβ_42_ surviving flies, whereas more than 30% of the w^1118^ flies and more than 15% of the Aβ_42_;Tip60 flies were remained alive (Fig. 3C). These results demonstrate that increasing Tip60 levels can protect against the decreased longevity of aged Aβ_42_ flies (Fig. 3C).

### Tip60/HDAC2 balance is disrupted during early Aβ_42_-induced neurodegeneration

Prior studies have shown inappropriately increased levels of HDAC2 in the brain of an AD-associated CK-p25 mouse model that is conserved in the hippocampus of AD patients (Graff et al. 2012). More recently, we demonstrated that disruption of Tip60/HDAC2 homeostasis consisting of increased HDAC2 levels and decreased Tip60 levels is an early event in the brain of an AD-associated APP fly model that can be protected against by increased Tip60 levels. Nevertheless, whether this epigenetic disruption mechanism can be induced solely in response to Aβ_42_ production during early and late stages of neurodegeneration progression and whether Tip60 can protect against these alterations remained to be characterized. Thus, we asked whether levels of the *Drosophila* HDAC1/2 ortholog termed Rpd3 and Tip60 are altered in response to early and late Aβ_42_-induced neurodegeneration. As HDAC2 and Rpd3 play conserved roles in memory formation and synaptic activity in mouse and *Drosophila*, respectively (Guan et al. 2009; Tea et al. 2010; Fitzsimons and Scott 2011), we refer to Rpd3 here as an HDAC2 orthologue. We used Western blot analysis on bulk proteins isolated from the heads of staged third instar larvae and from the heads of aged 28-day flies expressing either Aβ_42_ or Aβ_42_;Tip60 under the control of the pan-neuronal elav-GAL4 driver. These results revealed a significant increase in HDAC2 protein levels during early larval Aβ_42_-induced neurodegeneration that persists into the aged 28-day adult brain and is protected against in both early and late neurodegenerative stages by increasing Tip60 levels (Fig. 4Ai, 4Bi, 4Aii, and 4Bii). Tip60 preferential acetylated lysine residues (Anamika et al. 2010) H4K16 (Renaud et al. 2016) and H4K12 (Wee et al. 2014) levels were also significantly reduced throughout Aβ_42_ neurodegeneration progression, and these alterations were protected against (Fig. 4Aiii, 4Biii, and Supplemental Fig. S4) by increased Tip60 levels (Fig. 4Ai and 4Bi). Together these results demonstrate that Aβ_42_ induction alone has the capacity to alter epigenetic HDAC2/Tip60 balance before amyloid plaque accumulation in the brain and that such epigenetic dysfunction is protected against by an increase in Tip60 levels throughout neurodegenerative progression.

**Figure 4.**
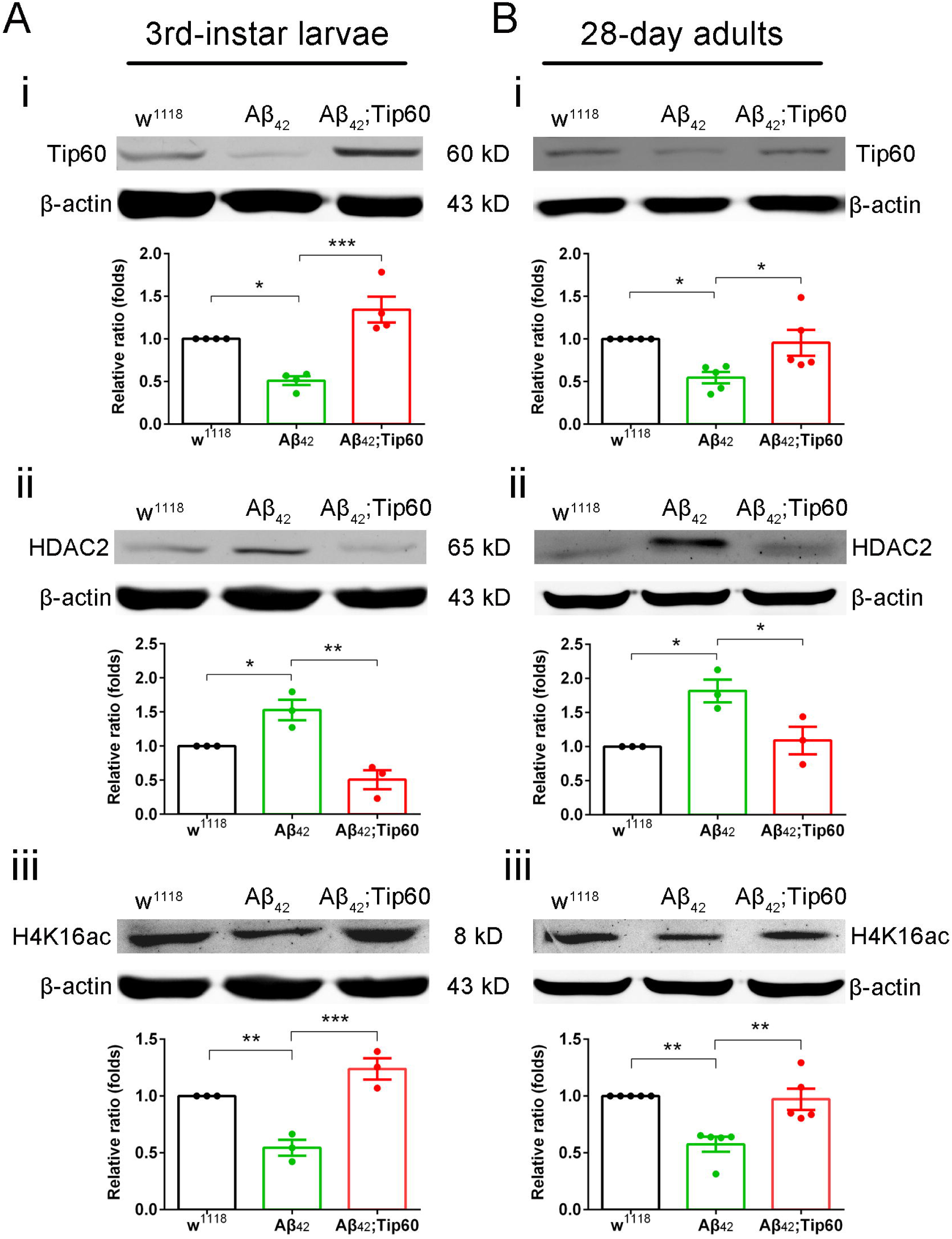
Increased Tip60 levels in the Aβ_42_ brain protects against disruption of Tip60/HDAC2 balance and concomitant reduction in histone acetylation levels. (A) Western blot using 3rd-instar larval heads shows early-stage changes. (i) Tip60 protein levels are decreased in the Aβ_42_ larval brain, and restored by Tip60 overexpression. n = 4. (ii) HDAC2 protein levels are increased in the Aβ_42_ larval brain, which is rescued by Tip60 overexpression. n = 3. (iii) Acetyl-H4K16 (H4K16ac) levels are decreased in the Aβ_42_ larval brain, which is restored by Tip60 overexpression. n = 3. (B) Western blot using 28-day adult heads shows late-stage changes. (i) Tip60 protein levels are decreased in the Aβ_42_ adult brain, which is restored by Tip60 overexpression. n = 5. (ii) HDAC2 protein levels are increased in the Aβ_42_ adult brain, which is rescued by Tip60 overexpression. n = 3. (iii) H4K16ac levels are decreased in the Aβ_42_ larval brain, which is restored by Tip60 overexpression. n = 5. Each biological repeat has 60 ∼ 70 larval or adult heads. \**p* < 0.05, \*\**p* < 0.01, \*\*\**p* < 0.001; one-way ANOVA with Tukey’s multiple comparisons test. All data are shown as mean ± s.e.m.

### Tip60 protects against early and late Aβ_42_-induced transcriptome-wide changes *via* different mechanisms

Evidence from gene expression (Grothe et al. 2018; Patel et al. 2019) and genetic variation (Karch et al. 2014; Kunkle et al. 2019) studies indicate that alteration in gene control may contribute to AD. Nevertheless, whether gene alterations are induced solely by Aβ_42_ production during early and late stages of AD progression and whether Tip60 can protect against them remains to be determined as neurodegenerative gene studies predominantly focus on aged brain samples. To address these questions, we profiled transcriptional changes during early neurodegeneration stages modeled in Aβ_42_ larval brains and late AD stages modeled in the aged 28-day old Aβ_42_ fly brain. Given our finding that Tip60 protects against Aβ_42_-induced Tip60/HDAC2 disruption and cellular defects in the brain, we also asked whether increased levels of Tip60 would protect against potential gene alterations throughout early and late staged Aβ_42_-induced neurodegeneration.

For transcriptome analysis, RNA was isolated from the brains of staged third instar larval heads and from the heads of aged 28-day flies expressing either Aβ_42_ or Aβ_42_;Tip60 under the control of the pan-neuronal elav-GAL4 driver. We used RNA sequencing to quantify gene expression changes for approximately 13500 genes in the larval brain and 13700 genes in the aged adult brain (Supplemental Fig. S5 and Supplemental Tbl. S1). Analysis in the Aβ_42_ larval brain reveals a large proportion of misregulated genes that include 1480 upregulated genes and 1687 downregulated genes. These gene changes depicted by Volcano plot (Fig. 5Ai and 5Aii) and Venn diagram (Fig. 5Bi) analysis reveal a larger proportion of genes are inappropriately repressed during early stages of Aβ_42_-induced neurodegeneration and that this inappropriate repression in gene expression profile trends (10.94% of the total upregulated vs. 12.47% of the total downregulated) is protected against by increased Tip60 levels in the Tip60; Aβ_42_ larval brain. Similarly, in the aged 28-day adult flies, we observed the same trend of overall gene misregulation (Fig. 5Aiii) with a slightly larger proportion of genes being inappropriately downregulated (78 genes upregulated, 0.57% of the total vs. 81 genes downregulated, 0.59% of the total) in the aged Aβ_42_ fly brain (Fig. 5Aiii and 5Bii). However, in direct contrast to the Aβ_42_ larval brain, increasing Tip60 levels does not significantly increase the number of upregulated genes but substantially decreases the amount of upregulated genes (Fig. 5Aiv and 5Bii).

**Figure 5.**
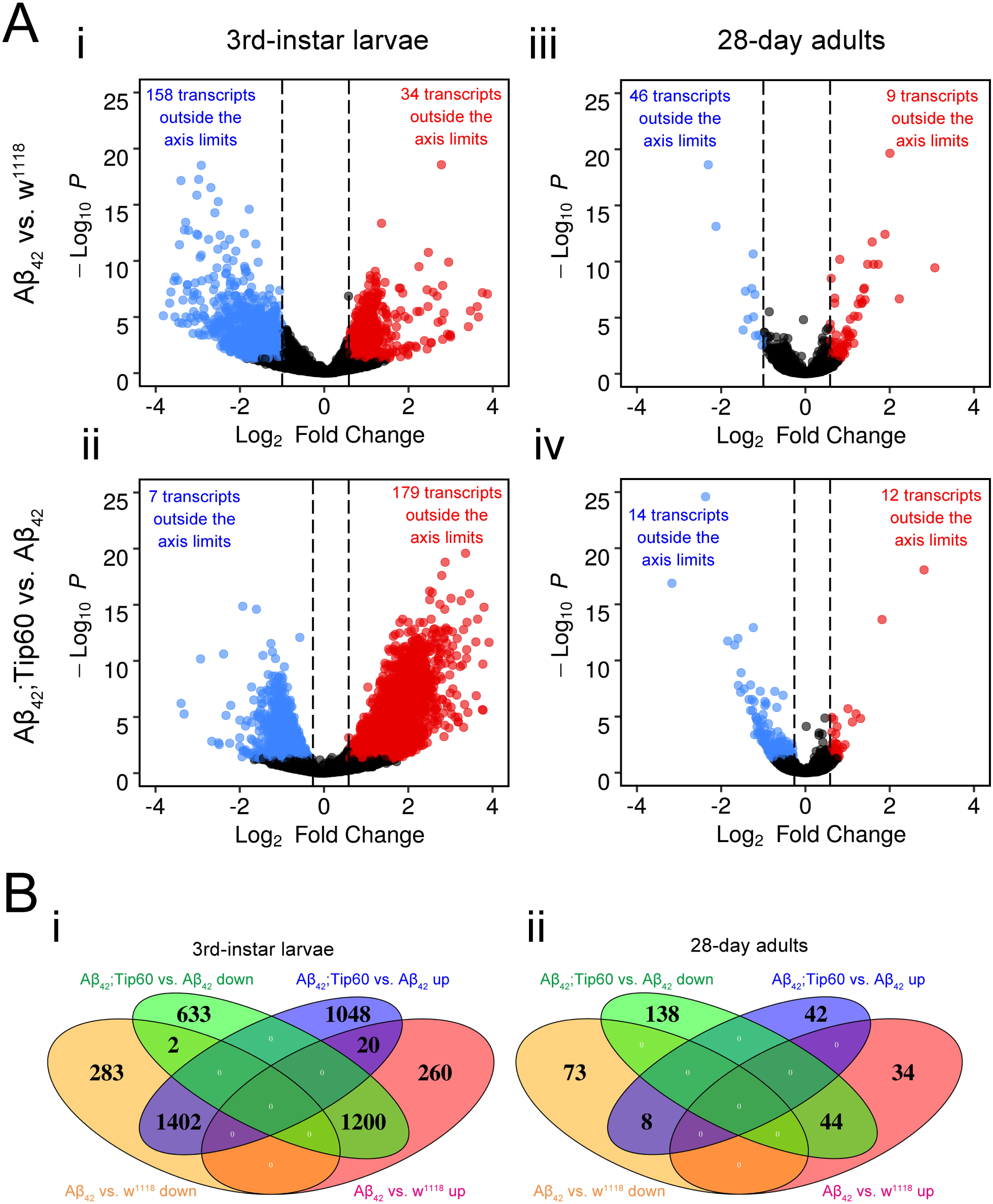
RNA-Seq reveals aberrant gene expression in the Aβ_42_ fly brain during early and late stage AD pathology that is rescued by increasing Tip60. (A) Volcano plots. (i) The differential gene expression in Aβ_42_ vs. w^1118^ in the 3rd-instar larval brain. (ii) The differential gene expression in Aβ_42_;Tip60 vs. Aβ_42_ in the 3rd-instar larval brain. (iii) The differential gene expression in Aβ_42_ vs. w^1118^ in the 28-day adult brain (many downregulated transcripts are outside the axis limits). (iv) The differential gene expression in Aβ_42_;Tip60 vs. Aβ_42_ in the 28-day adult brain. The RNAs falling into the upper left (blue dots) and upper right (red dots) quadrants are considered significantly altered. (B) Venn diagrams show that a high proportion of genes responding to Aβ_42_ display an effectively reversed differential expression when Tip60 levels are restored. (i) Distributions of differentially expressed RNAs between Aβ_42_ vs. w^1118^ and Aβ_42_;Tip60 vs. Aβ_42_ in 3rd-instar larvae. (ii) Distributions of differentially expressed RNAs between Aβ_42_ vs. w^1118^ and Aβ_42_;Tip60 vs. Aβ_42_ in 28-day adults. The intersection refers to the same genes between different comparisons. n = 2 ∼ 3 for larvae. Each biological repeat uses 30 ∼ 35 larval brains. n = 3 for adults. Each biological repeat uses 60 ∼ 70 adult heads.

To identify the biological processes that were altered under Aβ_42_-induced conditions and restored by increased Tip60 levels, functional annotation clustering was performed on the misregulated genes using gene ontology analysis (GSEA). The top 60 most significantly altered processes were compiled and classified under general umbrella terms (Fig. 6A and Supplemental Tbl. S2). They were next further categorized into transient (larval fly brain only), late-onset (aged 28-day fly brain only), and persistent (both early and late stages) expression classes (Fig. 6A and Supplemental Tbl. S3). Venn Diagram analysis of these biological process umbrella terms reveals well-established AD-associated processes that are repressed during early neurodgeneration progression modeled in Aβ_42_ larval brains such as serine peptidase activity, which plays a role in synaptic function and behavior (Almonte and Sweatt 2011) and general metabolic processes. Strikingly, the specific biological processes heat map analysis reveals a substantial number of gene subsets that are protected against by increased Tip60 levels (Fig. 6B). Of note, these processes predominantly include DNA packaging, RNA splicing, and RNA modifications that converge under the general function of transcriptional regulation, consistent with the role of Tip60 in gene control. Moreover, a substantial number of inappropriately upregulated gene processes are protected against by Tip60. Importantly, many of these processes are critical for neural function (axon guidance and axonogenesis, 8 processes/top 60, neuronal differentiation, 3 processes/top 60) and cell cycle control (cell division, 13 processes/top 60, cell cycle, 7 processes/top 60), consistent with changes previously associated with AD (Grothe et al. 2018) and with the well-characterized role for Tip60 in neural function and cell proliferation (Lorbeck et al. 2011). In direct contrast, in the aged 28-day adult Aβ_42_ brain, while a substantial number of inappropriate up and downregulated cellular processes were identified, increased levels of Tip60 protect only against the upregulation of helicase cellular processes. Intriguingly, detailed analysis of the cellular processes upregulated in response to increased Tip60 levels in the aged Aβ_42_ brain reveals significant enrichment in synaptic plasticity functions that include processes such as voltage-gated ion channel activity and neuronal projection function (Fig. 6C). Taken together, these results suggest that Tip60 mediates neuroprotection in early and late Aβ_42_-induced transcriptome-wide changes *via* different mechanisms with greater Tip60 specificity against Aβ_42_ changes during earlier stages of progression.

**Figure 6.**
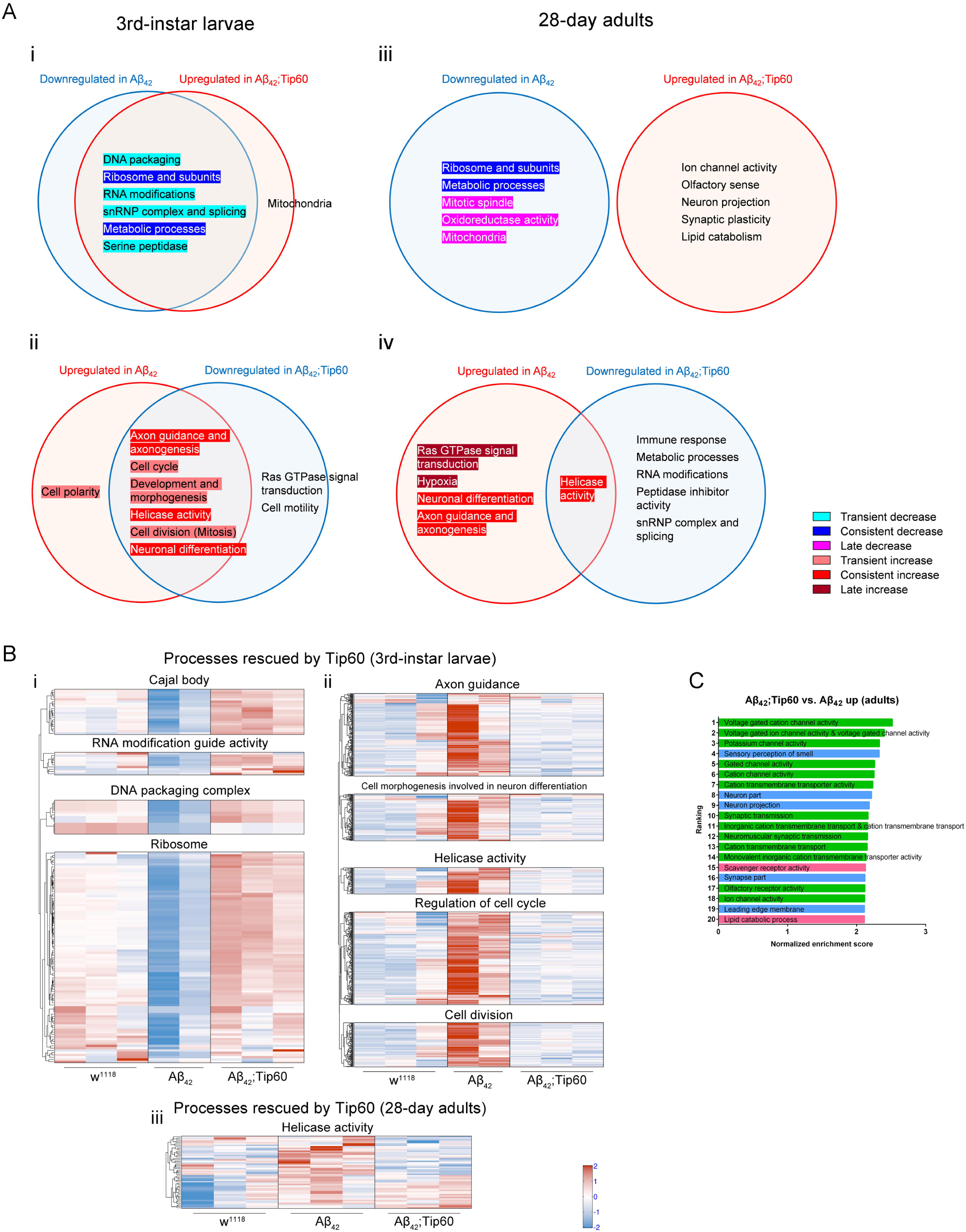
Gene Ontology (GO) analysis reveals different modes of action for Tip60 neuroprotection during early and late stages of Aβ_42_-induced neurodegeneration. (A) Venn diagrams show GO term associated enrichment in each of the significant functional categories. (i) Early and (iii) late Aβ_42_ vs. w^1118^ downregulated processes and Aβ_42_;Tip60 vs. Aβ_42_ upregulated processes. (ii) Early and (iv) late Aβ_42_ vs. w^1118^ upregulated processes and Aβ_42_;Tip60 vs. Aβ_42_ downregulated processes. (B) Heatmaps showing subsets of genes that are significantly misregulated by Aβ_42_ and rescued by Tip60. The most representative sub-processes which have the most significant gene size and lowest FDR q value under umbrella terms are selected for making this GO heatmap. (i) Heatmap of specific processes that are downregulated by Aβ_42_ and rescued by Tip60 in 3rd-instar larvae. (ii) Heatmap of specific processes that are upregulated by Aβ_42_ and rescued by Tip60 in 3rd-instar larvae. (ii) Heatmap of a specific process under the umbrella term of helicase activity that is upregulated by Aβ_42_ and rescued by Tip60 in 28-day adults. (C) Histogram of top 20 upregulated GO terms in adult fly Aβ_42_;Tip60 versus Aβ_42_ genotypes. These processes encompass functional neuroplasticity (green columns), structural neuroplasticity (blue columns), and lipid metabolism (pink columns). n = 2 ∼ 3 for larvae. Each biological repeat uses 30 ∼ 35 larval brains. n = 3 for adults. Each biological repeat uses 60 ∼ 70 adult heads.

## DISCUSSION

Here we report the first transcriptome-wide study assessing gene changes that arise during early and late stages of AD-associated neurodegeneration induced solely by induction of the neurotoxic human Aβ_42_ peptide in the *Drosophila* brain. A major novel finding from this study is that disruption of Tip60 HAT/HDAC2 balance involving increased HDAC2 and decreased Tip60 levels and robust gene expression alterations are induced solely by Aβ_42_. Strikingly, these alterations are an early event in neurodegeneration progression that arises several weeks before amyloid plaque accumulation is observed in the brain. Further, we found that substantially more genes are misregulated during early stages of Aβ_42_-induced neurodegeneration modeled in the larval brain (3167 genes) when compared to the brains from 28-day old aged adult flies (159 genes). While such early and robust dysregulation of epigenetic gene control was initially surprising, our findings can be explained by studies elucidating the biological function of Aβ_42_ protein. Aβ_42_ is a normal cellular metabolism product derived from the APP. Newly generated Aβ_42_ can be either released into the extracellular space or associate with plasma membrane lipid rafts that favors their accumulation into amyloid plaques. Studies show that the secreted soluble oligomeric and dimer prefibrillar species of the Aβ_42_ peptide can bind to multiple types of cell receptors and transduce neurotoxic signals into neurons causing cellular defects that include mitochondrial dysfunction, oxidative stress and transcriptional dysregulation (Chen et al. 2017). Inappropriate activation of signal transduction pathways can also arise via Aβ-mediated competition for binding of essential ligands to receptors. For example, soluble Aβ dimers cause glutamate excitotoxicity via blockage of glutamate reuptake in the synaptic cleft (Shankar et al. 2008; Lin et al. 2019; Zott et al. 2019) that activates multiple types of glutamate receptors and downstream cell signaling transduction cascades, pathologically altering gene expression profiles. Notably, our RNA-Seq results show that the topmost significant early gene alterations include those involved in metabolic cellular processes and neuronal function. Our findings are consistent with previously documented gene alterations associated with human AD pathology that have also been reported to detrimentally change in response to soluble Aβ-induced inappropriate cell signaling events (Grothe et al. 2018). Taken together, our finding that Aβ_42_ triggers more severe epigenetic gene dysregulation during the early stages of neurodegeneration functionally supports the concept that early-stage soluble Aβ dimers and oligomers (Chen et al. 2017) elicit enhanced cell surface receptor-mediated signal transduction effects over aged dependent insoluble Aβ_42_. Such Aβ plaque independent gene alterations can ultimately contribute significantly to AD pathologies.

Our study illustrates the power of *Drosophila* for the study of human AD neurodegeneration progression and specifically for elucidating alterations that arise during the early stages of the disease before Aβ_42_ plaque accumulation. Consistent with studies in CK-p25 AD-associated mouse model (Gjoneska et al. 2015), our transcriptome studies revealed gene alterations that we classified as transient (larval stage only), late-onset (28-day aged fly only), and constant (both). These coordinated alterations in biological processes are consistent with AD pathophysiology and likely reflect changes in both cell-type-specific expression profiles and cell types. Importantly, constant gene expression changes strongly feature neuroinflammation activation and metabolic inhibition that are key processes associated with AD pathology. Notably, we found the top 60 biologically processes upregulated by Aβ_42_ early neurodegenerative progression in the larval brain are predominantly neuronal processes related to neuroinflammation (Fig. 6Aii). For example, axon guidance processes upregulated in our studies and in human AD have recently been implicated in mediating immune and inflammatory responses in the postnatal period (Lee et al. 2019). The increased neuronal differentiation processes we observed can also be detrimental, leading to increased neurogenic-to-gliogenic fate switch (Paavilainen et al. 2018; Satir et al. 2020) and increased APP expression (Satir et al. 2020). Such gene alterations in early AD progression likely contribute to the learning and memory deficits observed in Aβ_42_ larvae (Fig. 3Ai and Supplemental Fig. S3Aii).

The majority of the top 60 processes downregulated by Aβ_42_ in the larval brain (Fig. 6Ai) are associated with gene expression regulation, consistent with the substantial gene expression alterations we observe during early AD stages in the larval brain. In the aged adult fly brain, the processes upregulated by Aβ_42_ are predominantly Ras superfamily of small GTPases as well as the constant inappropriate upregulation in neuronal processes. The small GTPases play crucial roles in neurogenesis, cell differentiation, gene expression, and synaptic plasticity, and have been implicated in AD pathogenesis (Qu et al. 2019). The top 60 downregulated processes in Aβ_42_ adults feature persistent changes in metabolic associated gene expression (Fig. 6Aiii) resulting in cellular respiration defects and compromised neural health (Fig. 6Aiv). Importantly, while our study uses transgenic flies overexpressing Aβ_42_ which likely display accelerated AD progression, we still observed many conserved cellular reactions that are hallmarks of human AD (De Strooper and Karran 2016) supporting the power of animal AD model systems like *Drosophila* for the study of human neurodegenerative disease.

Our results have therapeutic implications for AD as revealed by our finding that Tip60 protects against Aβ_42_-induced transcriptome-wide alterations *via* distinct mechanisms during early and state stages of neurodegeneration. During early neurodegeneration, Tip60 primarily protects biological processes that are specifically altered by Aβ_42_ that include gene regulatory, neuronal, and cell cycle processes (Fig. 5Ai and 5Aii), consistent with the role for Tip60 in mediating these functions. In direct contrast, during later neurodegenerative stages, Tip60-mediated gene protection under Aβ_42_ conditions become predominantly non-specific in that there are almost no processes altered by Aβ_42_ induction that are restored by Tip60. Rather, increased Tip60 in the Aβ_42_ aged brain promotes upregulation of numerous biological processes that are strikingly enriched for neural functions that include synaptic transmission, neural outgrowth, and multiple ion channel activity processes. Our findings support the following working model (Fig. 7). It has been well documented that well before amyloid plaque accumulation and massive neuronal cell death, neural function can become compromised in response to soluble Aβ-induced inappropriate cell signaling events (De Strooper and Karran 2016; Chen et al. 2017). We speculate that during these early stages, Tip60 cellular machinery remains functional, enabling increased Tip60 levels to compete with the factors that induce soluble Aβ_42_-induced gene alterations and specifically protect against them. However, upon disease progression, such specific Tip60 mediated neuroprotection mechanisms fail with overwhelming levels of Aβ accumulation and concomitant neuronal cell death. We speculate that Tip60-mediated restoration of neural function processes during these later stages of neurodegeneration represents a residual mechanism for surviving neurons to restore neuronal homeostasis and promote neural gene expression programs. Together, our findings demonstrate the therapeutic potential for Tip60 in both early and advanced stages AD that may provide the means for earlier and more selective treatments designed to restore neuronal histone acetylation homeostasis and function.

**Figure 7.**
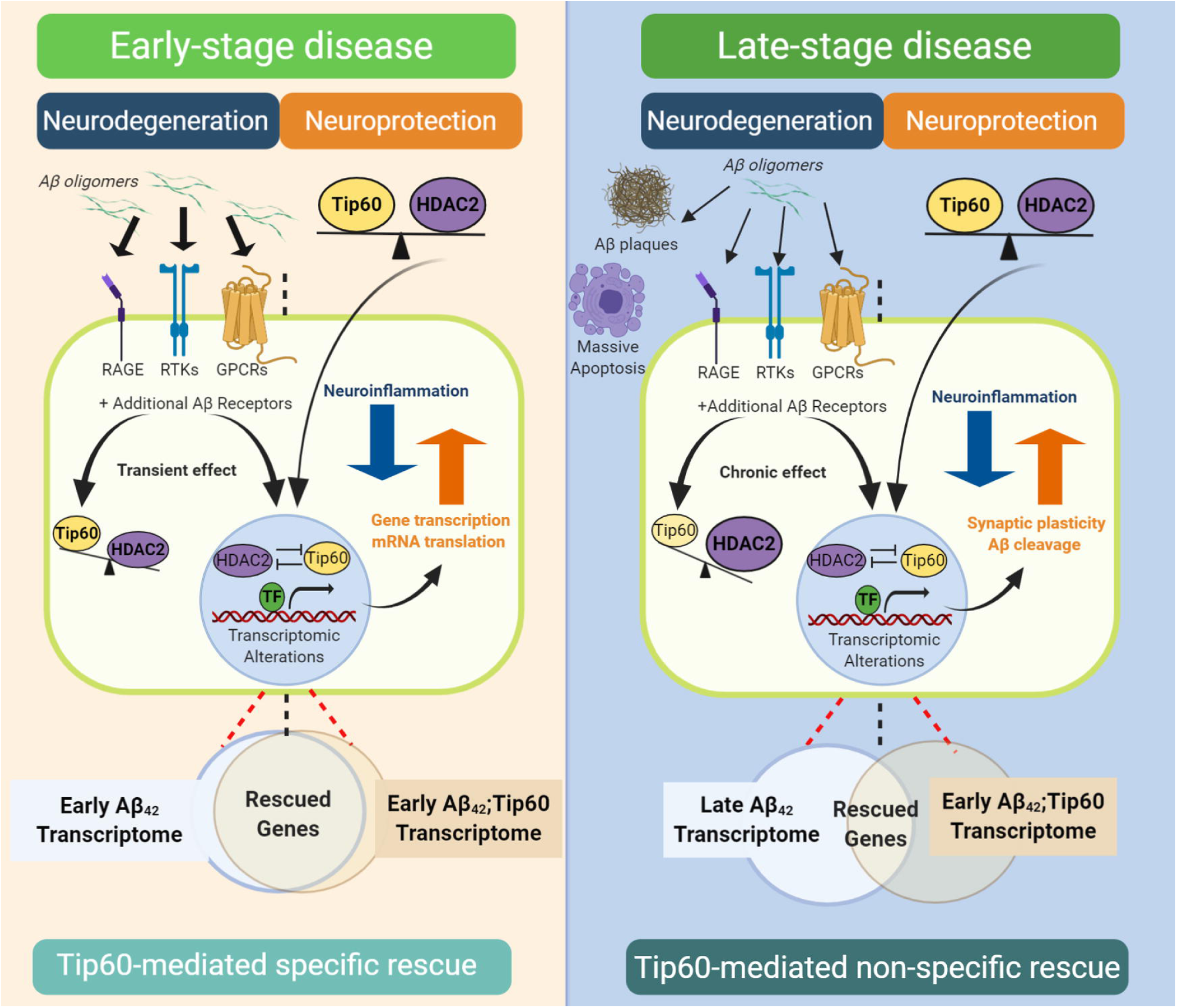
Working model for distinct modes of gene alterations and Tip60 neuroprotection during early and late stages of AD. Well before Aβ_42_-induced amyloid plaque accumulation and massive neural cell death, neural function becomes compromised in response to soluble Aβ-induced inappropriate cell signaling events that contribute to functional cognitive impairments. These soluble Aβ_42_ dimmers or oligomers can bind to multiple types of Aβ receptors to block appropriate ligand binding, leading to dysregulation of cellular signaling pathways that disrupt both Tip60 HAT/HDAC2 homeostasis and gene expression profiles. During these early stages, Tip60 cellular machinery remains stable and functional, enabling increased Tip60 levels to specifically compete with the factors that induce soluble Aβ_42_-induced gene alterations and protect against them. However, during late staged neurodegeneration, we speculate that while increased Tip60 levels can still promote induction of neuronal genes, Tip60 mediated protection against specific Aβ_42_ induced gene alterations is disrupted due to general cellular machinery failure. Abbreviations: receptor for advanced glycation endproducts, RAGE; receptor tyrosine kinases, RTKs; G-protein-coupled receptors, GPCRs; transcription factor, TF.

## MATERIALS AND METHODS

### Fly stocks

All fly lines were raised under standard conditions at 20-25LJ on standard yeasted *Drosophila* media (Applied Scientific Jazz Mix *Drosophila* Food; Thermo Fisher Scientific). The pan-neuronal driver elav-Gal4^C155^ (#458) and the transgenic UAS line carrying human Aβ_42_ (#33769) were purchased from Bloomington *Drosophila* Stock Center. We generated stocks carrying UAS-Tip60 (UAS-dTip60^WT^ C) previously (Lorbeck et al. 2011). The transgenic UAS-Tip60 fly line was crossed into the UAS-Aβ_42_ background using standard genetic techniques, as previously described (Pirooznia et al. 2012), to generate the UAS-Aβ_42_;UAS-Tip60 double transgenic fly line. The w^1118^ line (Bloomington *Drosophila* Stock Center, #3605) served as the genetic background control. All experimental crosses were performed at an average physiological temperature of 20-25LJ.

### Immunofluorescence, imaging, and quantification

For anti-Aβ_42_ immunofluorescence, larval or adult brains were dissected in PBS, fixed in fixation buffer containing 0.7% paraformaldehyde and 0.9% lysine for 1 hour at room temperature, washed three times in PBS containing 0.5% Triton X-100 (PBST) for 15 minutes each time at room temperature, and blocked for 1 hour at room temperature in PBST containing 5% normal goat serum, and incubated with primary anti-Aβ42 (1:100, Millipore, #05-831-I) antibody in blocking solution overnight at 4LJ. Samples were washed three times in PBST for 15 minutes each time at room temperature and incubated with goat anti-mouse Alexa Fluor 488 (1:300, Invitrogen, #A28175) and propidium iodide (PI, a final concentration of 1.5 μM) for 2 hours at room temperature. After washing three times in PBST for 15 minutes each time, samples were mounted in VECTASHIELD Antifade Mounting Media (VECTOR LABORATORIES).

TUNEL staining was performed according to the manufacturer’s instructions (*In Situ* Cell Death Detection Kit, Fluorescein, Roche, #11 684 795 910). In brief, brain fixation, blocking, rinsing, and PI nuclear staining procedures are the same as anti-Aβ_42_ immunofluorescence. The only difference is instead of incubating the brains in the primary antibody, incubate the brains in the TUNEL reaction mixture in the humidified atmosphere for 1 hour at 37LJ in the dark. TUNEL positive control experiment was performed using DNase LJ (ZYMO RESEARCH, #E1010)-treated brains.

Confocal microscopy was performed using a ZEISS microscope (LSM 700, ZEISS United States). The optical intervals were 5.94 μm z-sections for 100× magnifications and 0.79 μm z-sections for 200× magnifications. The optical intervals were determined by the optimized pinhole diameters which are 33.3 μm at 1 Airy Unit (AU) for 100× magnification and 25.1 μm at 1 AU for 200× magnification. Consecutive z-stacks through the entire Kn were used for quantification. Consecutive subsets of the z-stacks approximately at the level of center Kn were used for the final projection and display. The quantification of Aβ plaques and apoptosis in different genotypes was measured under 200× magnification using Image J software.

### Western Blot

3rd-instar larval heads and 28-day adult heads were dissected for protein extraction. The protein was analyzed using a BCA assay (Thermo Scientific, #23225). Protein extracts (total protein: 30 μg) were electrophoresed on 10% sodium dodecyl sulfate-polyacrylamide gels and transferred to polyvinylidene difluoride membranes for small molecules (H4K12ac and H4K16ac) or nitrocellulose membranes for large molecules (Tip60 and Rpd3). The blots were blocked in TBST buffer containing 5% nonfat milk and then incubated overnight at 4°C with the primary rabbit polyclonal anti-Tip60 (1:1000, abcam, #ab23886), rabbit polyclonal anti-Rpd3 (1:1000, abcam, #ab1767) or mouse monoclonal anti-β-actin (1:300, DSHB, #JLA20) antibody diluted in TBST. The blots were then rinsed and incubated with the appropriate secondary antibodies (1:15000, IRDye 800CW Goat anti-Mouse, LI-COR, #925-32210, or 1:15000, IRDye 680RD Goat anti-Rabbit, LI-COR, #926-68071) for 1 hour at room temperature and scanned using the Western Blot detection system (Odyssey). The total proteins were normalized to β-actin. Densitometry was determined by band intensity, using Image J software.

#### RT-qPCR

3rd-instar larval brains or 28-day adult heads were homogenized, and total RNA was isolated using the QIAGEN RNeasy Mini Kit (QIAGEN, #74106) following the manufacturer’s protocol. cDNA was synthesized using random primers (Roche, #11034731001). Samples were amplified for 35 cycles using the 7500 Real-Time PCR system (Applied Biosystems). The RNAs were normalized to RPL32. The primer sequences are: Aβ sense: GCAGAATTCCGACATGACTCAG, anti-sense: GCCCACCATGAGTCCAATGA; Tip60 sense: CCTTCCACGACCTGAACTCC, anti-sense: CTCGGCCTGAGGCTTGTAAC; RPL32 sense: AGGGTATCGACAACAGAGTGC, anti-sense: CTTCTTGAATCCGGTGGG.

### RNA-sequencing (RNA-Seq)

3rd-instar larval brains or 28-day adult heads were homogenized, and total RNA was isolated using the QIAGEN RNeasy Mini Kit (QIAGEN, #74106) following the manufacturer’s protocol. Total RNA quantity, quality, and purity were determined using the Agilent 2100 bioanalyzer and Nanodrop spectrophotometer. Only the RNAs with an RNA integrity number (RIN) ≥ 6.0 were used for subsequent sequencing. 100 ng of total RNA was used to prepare libraries using TruSeq Stranded Total RNA kit (Illumina, CA, USA) following the manufacturer’s protocol. The final libraries at the concentration of 4 nM were sequenced on NextSeq 500 using 75 bp paired-end sequencing. Raw FASTQ sequencing reads were mapped against the reference genome of *Drosophila melanogaster* (Ensembl version BDGP6) using RNA-Seq by Expectation-Maximization (RSEM) (Li and Dewey 2011). Total read counts and normalized Transcripts Per Million (TPM) were obtained using RSEM’s calculate-expression function. Before, differential expression, batch effects, or sample heterogeneity was tested using iSeqQC (https://github.com/gkumar09/iSeqQC).

### RNA-Seq data analysis

Differential gene expression was tested using the DESeq2 package in R/Bioconductor (Love et al. 2014). Volcano plot, Venn diagram, and heatmaps were constructed using R/Bioconductor. To set up cutoff criteria in volcano plots and Venn diagrams, we took into consideration the biological significance and calculated the values accordingly. For Aβ_42_ vs. w^1118^, genes that were downregulated or upregulated by greater than or equal to 50% were considered to be differentially expressed in the Aβ_42_ fly brain. The corresponding parameters are adjusted *p* < 0.05, fold change (FC) of Aβ_42_ vs w^1118^ ≤ 0.5 (log_2_FC ≤ − 1), and *p* < 0.05, FC of Aβ_42_ vs w^1118^ ≥ 1.5 (log_2_FC ≥ 0.585), respectively. Additionally, genes restored to at least the average of w^1118^ and Aβ_42_ were considered to be partially rescued by Tip60 in the Aβ_42_;Tip60 fly strain. That is, FC of Aβ_42_;Tip60 vs. w^1118^ ≥ 0.75 and FC of Aβ_42_;Tip60 vs. w^1118^ ≤ 1.25, respectively. Accordingly, FC of Aβ_42_;Tip60 vs. Aβ_42_ ≥ 1.5 (log_2_FC ≥ 0.585) and FC of Aβ_42_;Tip60 vs. Aβ_42_ ≤ 0.833 (log_2_FC ≤ − 0.263) were calculated. Therefore, genes were considered differentially expressed if they had adjusted *p* < 0.05 and log_2_FC ≤ − 1 or log_2_FC ≥ 0.585 for the comparison of Aβ_42_ vs. w^1118^, or adjusted *p* < 0.05 and log_2_FC ≤ − 0.263 or log_2_FC ≥ 0.585 for the comparison of Aβ_42_;Tip60 vs. Aβ_42_. Gene ontology (GO) analysis was performed according to the Gene Set Enrichment Analysis (GSEA). To present GO processes without redundancies, we manually merged overlapping GO terms into umbrella terms according to the principles listed on the site http://wiki.geneontology.org/index.php/Merging_Ontology_Terms.

#### Larval olfactory associative memory assay

Larval olfactory associative memory assay was performed according to the well-established protocol (Honjo and Furukubo-Tokunaga 2005). Briefly, 3rd-instar larvae were conditioned on agar plates to learn to associate appetitive reinforcer sucrose and odorant. After training, larvae were transferred to the center of a new agar plate and given a choice to select between test odorant and control. The numbers of animals moving in the indicated semicircular areas were counted, and the response index (RI) is calculated accordingly by dividing the difference of the number of animals in odor and control by the total number of animals. Olfactory associative memory performances for indicated genotypes were plotted in ΔRI. ΔRI = RI (LIN/SUC) – RI (LIN/DW). To eliminate the effect of locomotor function on memory performance, we normalized the RIs of all of the genotypes using their respective moving speed. The moving speed was calculated by a Tracker software (http://physlets.org/tracker/) using the videos of distance individual larva traveled in one minute. Olfactory and gustatory control experiments were performed accordingly (Honjo and Furukubo-Tokunaga 2005).

#### Adult olfactory associative memory assay

Adult olfactory associative memory assay was performed as described (Malik and Hodge 2014) with minor revisions. The T-maze equipment was purchased from MazeEngineers (Boston, MA, USA). We trained the flies by electroshock consisting of twelve 3.75-second pulses with 1.25-second inter-pulse intervals paired with one odor (3-octanol, OCT, 1:100 or 4-methyl cyclohexanol, MCH, 1:67) for 60 seconds and subsequently exposed the flies to a second odor without electroshock for 60 seconds. Flies were tested following one session of training. Thirty minutes after training, STM is measured by allowing flies to choose between the two scents for 120 seconds. Olfactory associative memory performances for indicated genotypes were plotted in performance index (PI). PI was calculated by subtracting the number of flies making the incorrect choice from those making the correct one, divided by the total number of flies. The final PI was calculated by averaging the PI of the experiment in which OCT was the shock-paired odor and one in which MCH was the shock-paired odor. This averaged PI removes any bias of the flies having a higher performance for one odor. For olfactory control experiments, the absolute odor avoidance response was quantified by exposing naïve flies to each odor (OCT or MCH) or air in the T-maze. After 120 seconds, the numbers of flies in each arm of the T-maze were counted, and the PI was calculated for each odor individually. In the shock reactivity control experiments, the ability to sense and escape from electric shock was quantified by inserting electrifiable grids into both arms of the T-maze and delivering shock pulses to one arm. Flies were placed to the choice point of the T-maze, where they could choose between the two arms. After 120 seconds, the center compartment was closed, trapping flies in their respective divisions and the PI was calculated.

#### Larval and adult locomotion assays, and survival assay

For larval locomotion, we performed line crossing assay, righting assay, and body wall contraction assay, as described before (Mudher et al. 2004). Briefly, line crossing assay tests the number of lines the larval head crosses in 30 seconds. Righting assay tests the time it takes for the larva to turn from the ventral side up to the ventral side down. Body wall contraction assay tests the number of peristaltic contractions from back to the front in 30 seconds. For adult locomotion, we utilized the negative geotaxis assay (Krashes and Waddell 2008). Adult flies were transferred to an empty vial (9.3 centimeters length and 2.3 centimeters in diameter). The geotaxis index was determined by recording the percentage of flies to reach the upper vial in 10 seconds. For survival assay, a total of more than 100 flies were prepared for each genotype. The survival rate of the flies has been evaluated every two or three days, counting the number of live flies until all the flies are dead.

#### Statistical analyses

Student’s t-tests were used for comparison between parameters from two groups. One-way ANOVA tests with Tukey’s multiple comparison test were used for comparison between parameters from various groups. A log-rank test with multiple adjustments was used for survival assay. GraphPad Prism 6 (GraphPad Software, Inc.) was used for all statistical analyses.

## ACKNOWLEDGMENTS

Felice Elefant is supported by a National Institutes of Health (NIH) R01 grant (no. 5R01NS095799-04). Special thanks to Dr. James Hodge (University of Bristol) for providing adult fly memory assay (T-maze) expertise and Dr. Gaurav Kumar (Thomas Jefferson University) for RNA-Seq data analysis expertise. We thank the undergraduates of our lab, Ilayda Erkan, Sundus Pervez, Janine Yang, and Nazaarah Abdul-Aziz for their help with all the experiments. We would like to thank the Drexel University Cell Imaging Center for imaging facilities and assistance and Thomas Jefferson University RNA-Seq facility for sequencing assistance.

## AUTHOR CONTRIBUTIONS

F.E. and H.Z. conceived the project. H.Z. and S.M. generated the Aβ_42_;Tip60 fly line. H.Z. performed the immunofluorescence, RNA-Seq, and RT-qPCR. H.Z. and B.C.K. performed the Western blot. H.Z., T.V.R. and S.M. performed the behavioral assays. B.C.K., H.Z., and A.B. performed the RNA-Seq data analysis. E.M.A. helped with the RNA-Seq sample preparation. M.B. helped with the review and revision of the paper. H.Z. and F.E. wrote the article.

