## Supplemental material for "Tip60 protects against amyloid-β peptide-induced transcriptomic alterations via different modes of action in early versus late stages of neurodegenerative progression"

Running Title: Epigenetic gene changes during AD progression

**SUPPLEMENTAL MATERIAL LIST**

**Supplemental figures in this Word document**

Supplemental Figure S1. Validation of Aβ_42_ and Tip60 transgene expression.

Supplemental Figure S2. TUNEL positive controls.

Supplemental Figure S3. Behavioral control experiments.

Supplemental Figure S4. Additional Western blot of another Tip60-specific histone acetylation mark H4K12ac.

Supplemental Figure S5. Heatmaps of all the genes.

**Supplemental tables not included in this Word document**

Supplemental Table S1. Differential gene expression (Excel file).

Supplemental Table S2. GSEA (Excel file).

Supplemental Table S3. GSEA with further categorization (Excel file).

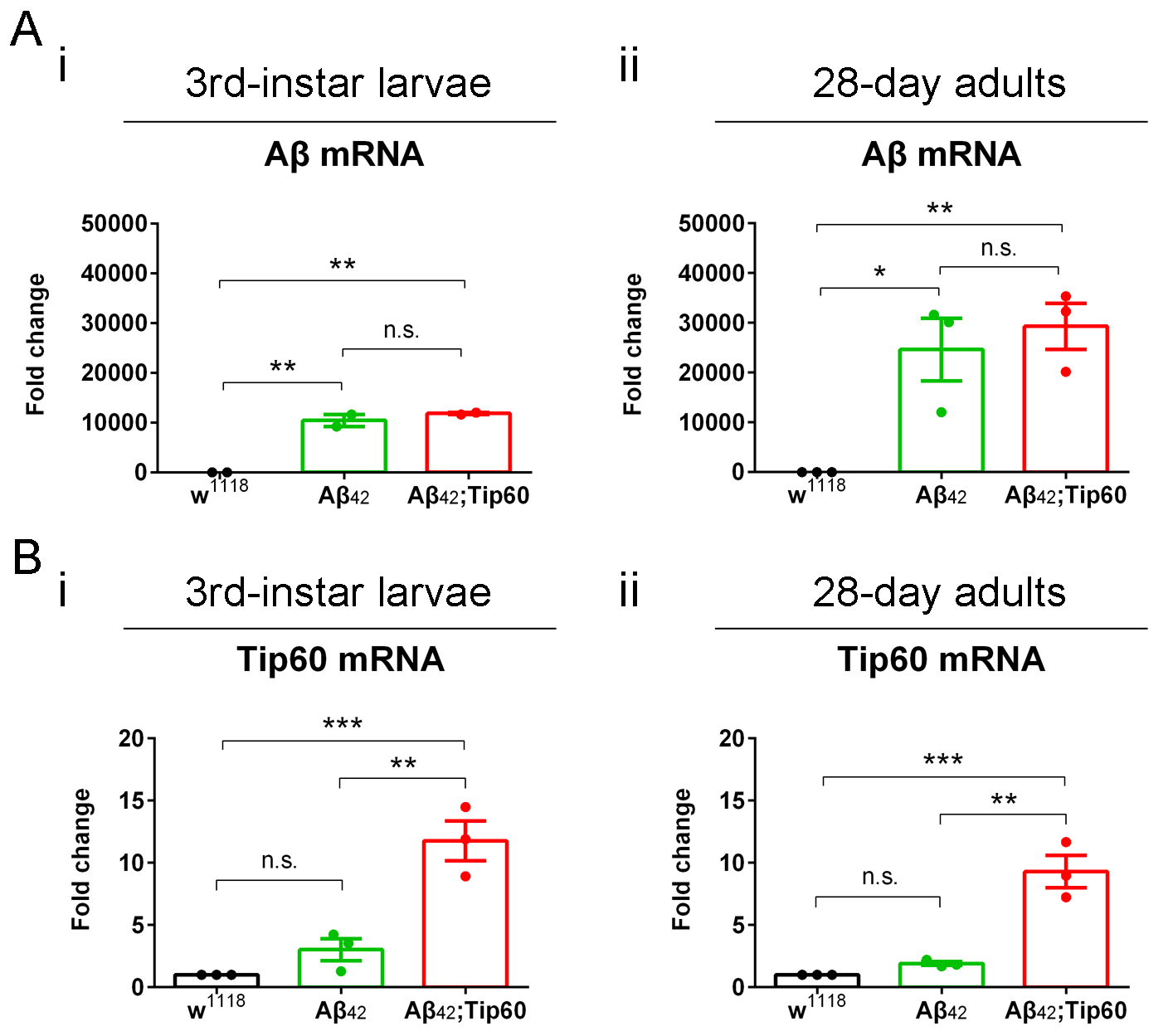

Supplemental Figure S1. qPCR validates Aβ_42_ and Tip60 transgene expression. (A) Negligible levels of Aβ_42_ mRNA are detected in w^1118^ control brains, and very high levels of Aβ_42_ mRNA are detected in both Aβ_42_ and Aβ_42_;Tip60 brains across early and late developmental stages. Aβ_42_ mRNA levels are comparable between Aβ_42_ and Aβ_42_;Tip60 brains. (i) RT-qPCR is performed using 3rd-instar larval brains. n = 2. (ii) RT-qPCR is performed using 28-day adult heads. n = 3. (B) Tip60 mRNA is effectively restored in the Aβ_42_;Tip60 fly brain early in life, and this restoration persists over time. (i) RT-qPCR is performed using 3rd-instar larval brains. n = 3. (ii) RT-qPCR is performed using 28-day adult heads. n = 3. Fold change is calculated using 2^-ΔΔCt^ method with RPL32 as the endogenous control. Each biological repeat uses 30 ~ 35 larval brains or 60 ~ 70 adult heads. **p* < 0.05, ***p* < 0.01, ****p* < 0.001; one-way ANOVA with Tukey’s multiple comparisons test. All data are shown as mean ± s.e.m.

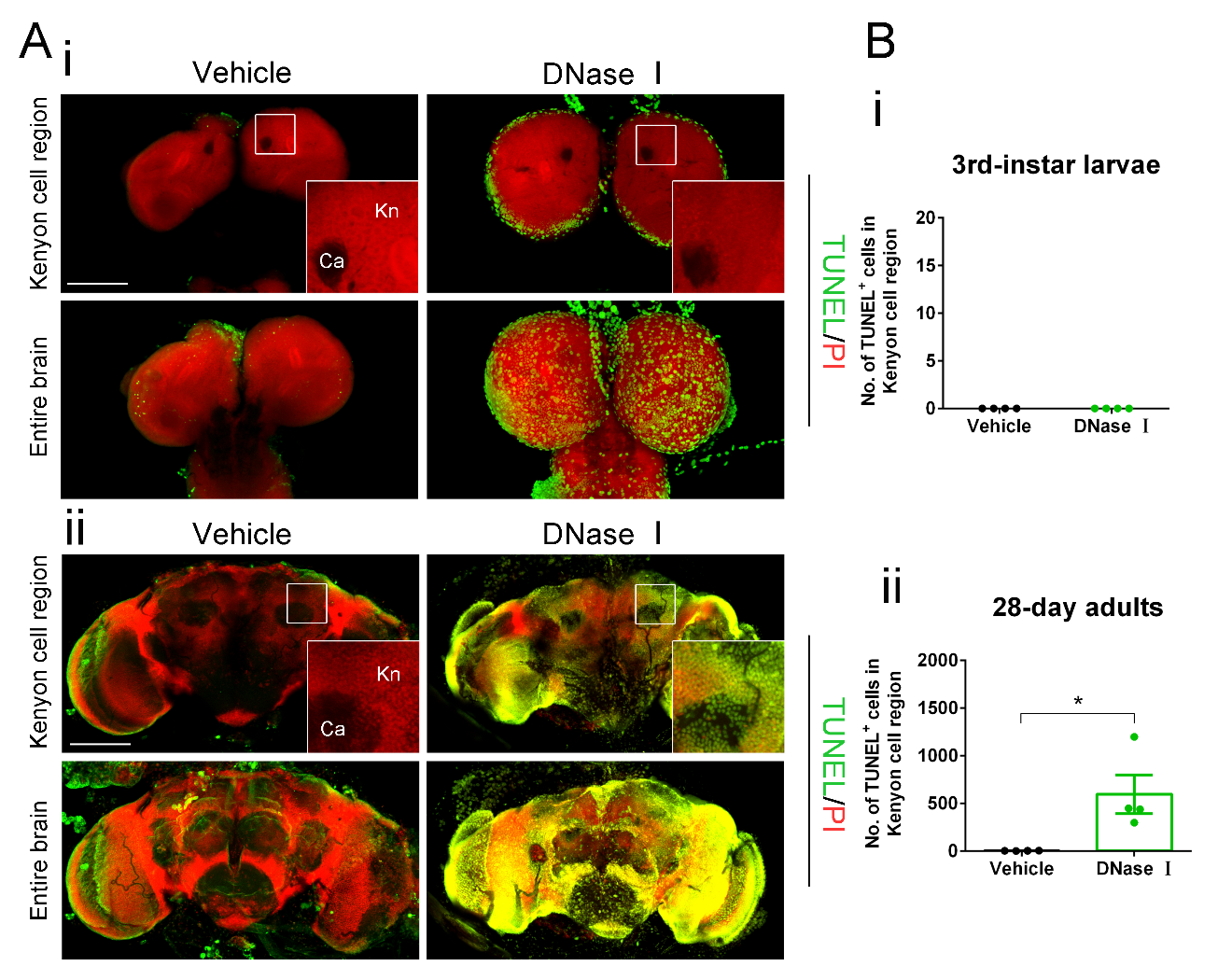

Supplemental Figure S2. The DNase Ⅰ-treated larval and adult brains used as TUNEL positive controls show massive apoptosis. (A) Representative confocal images of neuronal apoptosis visualized by TUNEL staining (green) of brains treated by vehicle or DNase Ⅰ are displayed. Nuclei were stained with PI (red). The Kenyon (Kn) cell region (boxed) is zoomed in to display Kenyon cells and apoptotic signals. (i) Immunostaining of DNase Ⅰ-treated 3rd-instar larval brain shows significantly more apoptotic signals than the vehicle-treated brain. (ii) Immunostaining of DNase Ⅰ-treated 28-day adult brain shows considerably more apoptotic signals than the vehicle processed brain. Scale bar represents 100 μm. (B) TUNEL signal was quantified by counting the number of TUNEL positive cells in the Kenyon cell region in larval (i) and adult (ii) brains. n = 4. **p* < 0.05; one-way ANOVA with Tukey’s multiple comparisons test. Data are shown as mean ± s.e.m.

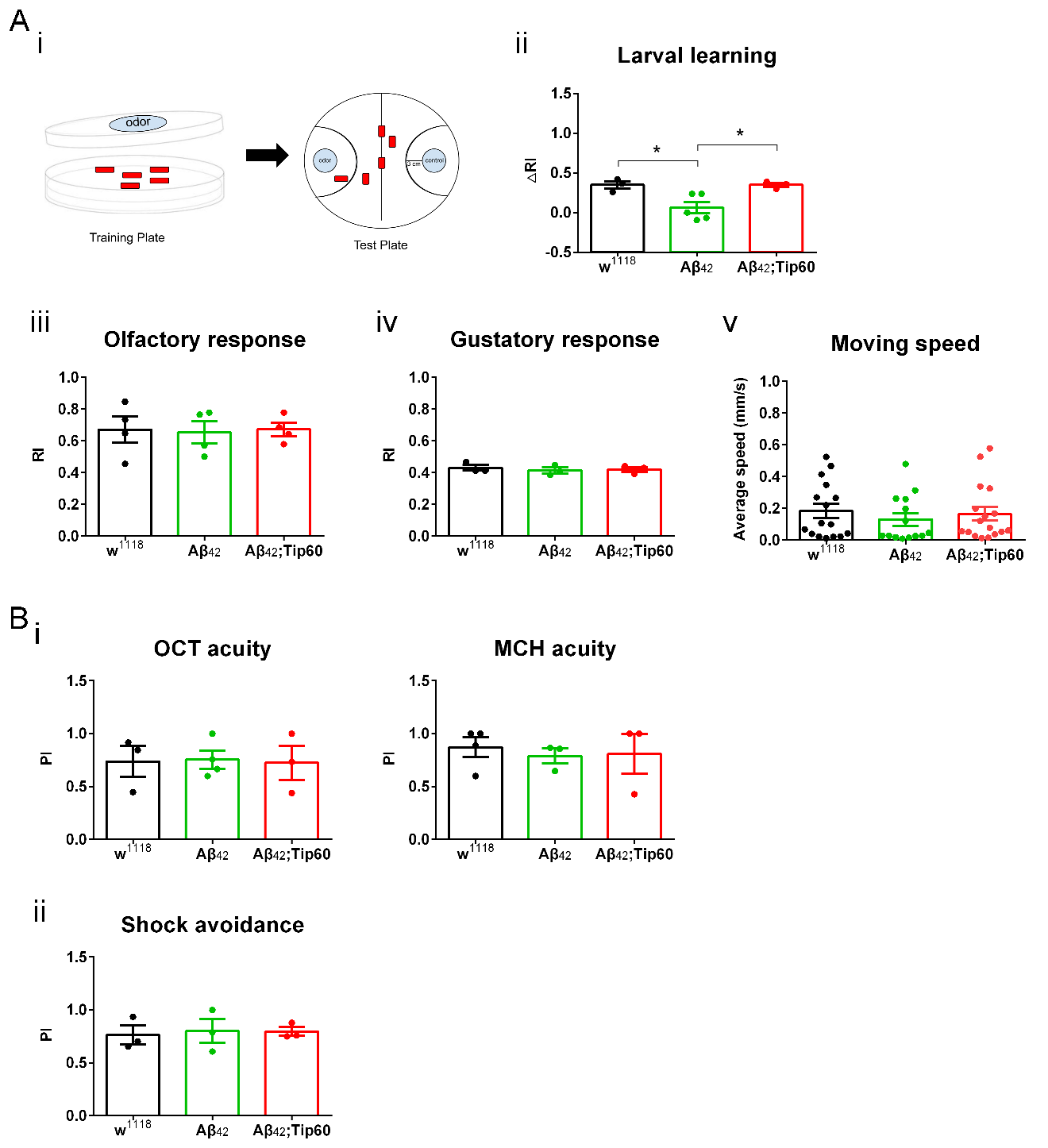

Supplemental Figure S3. Behavioral control experiments for larvae and adults. (A) Larval STM control experiments show learning ability, sensory acuity, and locomotion speed among all three genotypes. (i) A single odor paradigm for olfactory learning and memory. (ii) w^1118^ and Aβ_42_;Tip60 larvae show normal learning performance, whereas Aβ_42_ larvae only display a residue learning ability with a ∆RI close to zero. n = 3 ~ 5. each biological repeat uses 60 ~ 100 larvae. (iii) Larvae of different genotypes display comparable olfactory acuities to the odor linalool (LIN). n = 4. Each biological repeat uses about 30 larvae. (iv) Larvae of different genotypes display comparable gustatory acuities to the natural reinforcer sucrose. n = 3. Each biological repeat uses about 30 larvae. (v) Aβ_42_ larvae show a slight defect in moving speed compared to w^1118^ and Aβ_42_;Tip60 larvae, but this trend is not statistically significant. The locomotion speed of all the indicated genotypes was used to normalize ∆RIs in the learning and STM assays. n = 14 ~ 17. (B) Adult STM control experiments show similar sensory acuities and shock avoidance among all three genotypes. (i) Adults of different genotypes display comparable olfactory acuities to odor. n = 3 ~ 4. Each biological repeat uses at least 30 adults. (ii) Adults of different genotypes display comparable shock avoidance. n = 3. Each biological repeat has at least 30 adults. **p* < 0.05; one-way ANOVA with Tukey’s multiple comparisons test. All data are shown as mean ± s.e.m.

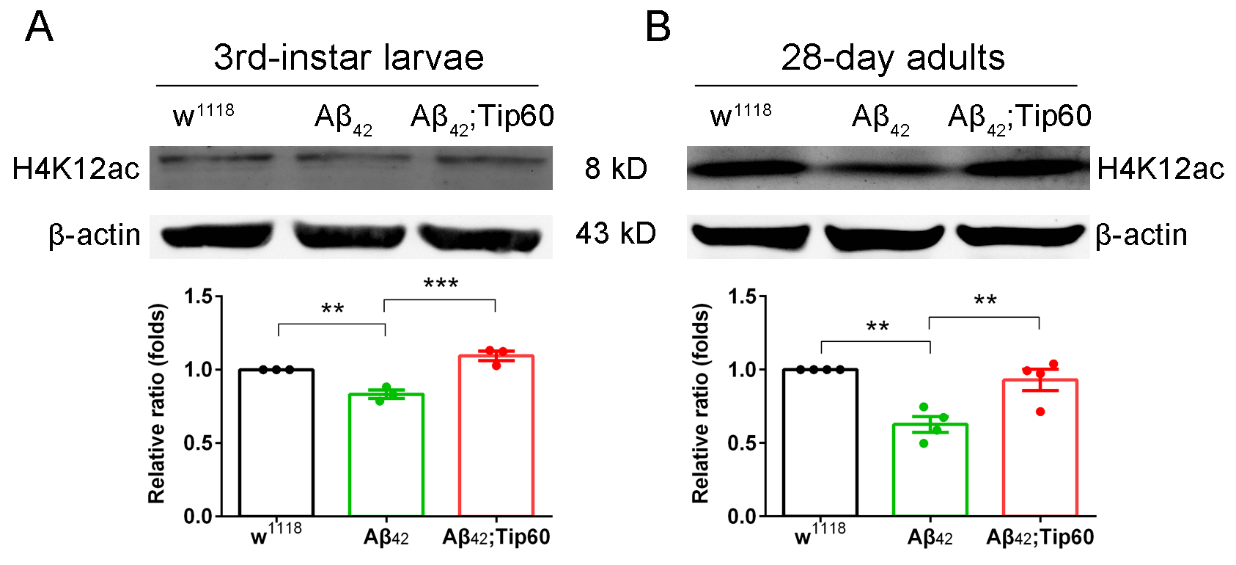

Supplemental Figure S4. Reduced Tip60-specific H4K12 acetylation (H4K12ac) in the Aβ_42_ fly brain is restored by Tip60 overexpression across the early and late disease stages. (A) Western blot is performed using 3rd-instar larval heads. n = 3. (B) Western blot is performed using 28-day adult heads. n = 4. Each biological repeat uses 60 ~ 70 larval or adult heads. ***p* < 0.01, ****p* < 0.001; one-way ANOVA with Tukey’s multiple comparisons test. Data are shown as mean ± s.e.m.

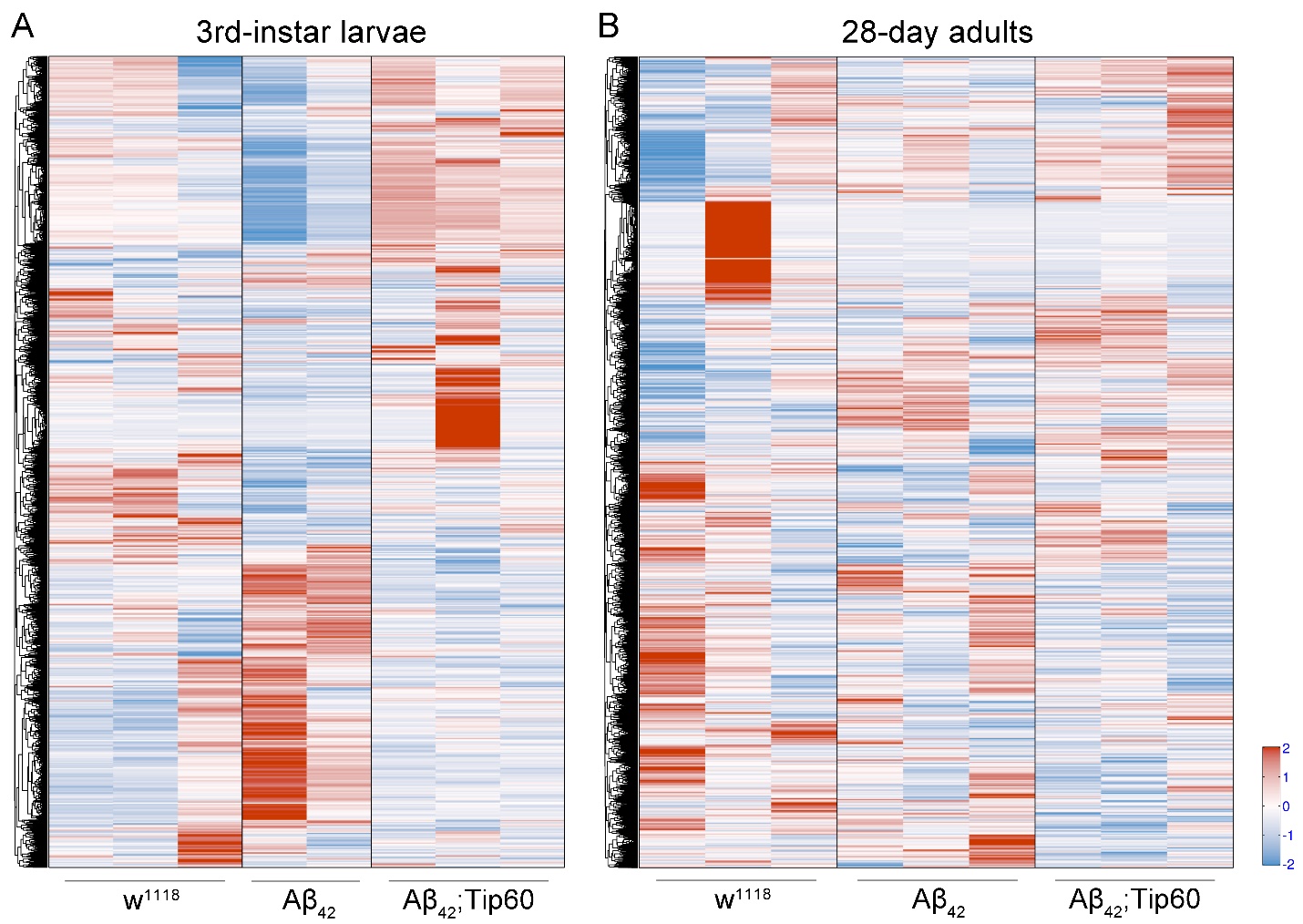
Supplemental Figure S5. Heatmaps showing differential gene expression. Compared with 28-day adults, the 3rd-instar larval gene expression profile displays a significantly more specific Tip60-mediated rescue of genes that are misregulated by Aβ_42_. Expression higher than the mean is displayed as shades of red, lower than the mean as shades of blue. (A) Heatmap of larval brain RNA-Seq. n = 2 ~ 3. Each biological repeat uses 30 ~ 35 larval brains. (B) Heatmap of adult head RNA-Seq. n = 3. Each biological repeat uses 60 ~ 70 adult heads.
